# Clustering NMR: Machine learning assistive rapid (pseudo) two-dimensional relaxometry mapping

**DOI:** 10.1101/2020.04.29.069195

**Authors:** Weng Kung Peng

## Abstract

Low-field nuclear magnetic resonance (NMR) relaxometry is an attractive approach for point-of-care testing medical diagnosis, industrial food science, and *in situ* oil-gas exploration. One of the problem however is, the inherently long relaxation time of the (liquid) sample, (and hence low signal-to-noise ratio) causes unnecessarily long repetition time. In this work, we present a new class of methodology for rapid and accurate object classification using NMR relaxometry with the aid of machine learning. We demonstrate that the sensitivity and specificity of the classification is substantially improved with higher order of (pseudo)-dimensionality (e.g., 2D or multidimensional). This new methodology (termed as Clustering NMR) is extremely useful for rapid and accurate object classification (in less than a minute) using the low-field NMR.

## Introduction

High resolution nuclear magnetic resonance (NMR) spectroscopy is a powerful and attractive approach in biochemistry (e.g., protein analysis^[1]^, metabolomics^[2–4]^) and inorganic chemistry^[5]^. In the recent years however, with the rapid advances in NMR engineering (e.g., IC-based spectrometer^[6–12]^, microfluidic-based chip^[13–17]^, artificial intelligence^[18,19]^) utilizing small foot-print permanent magnet, the time-domain NMR instrumentations have seen a myriad of interesting applications from point-of-care testing (PoCT) medical diagnosis^[7,20–23]^, industrial food science ^[24,25]^, and *in-situ* oil-gas exploration^[26,27]^.

Biochemical information is typically detected and encoded in the frequency domain (‘chemical shift’) in the high-field NMR. In contra, the low-field NMR, information is encoded in the time domain, with the dephasing of the spin-spin relaxation (T_2_ relaxation) of the water-proton of the observed sample used as diagnostic criterion^[20,21]^. Time domain NMR however, suffers from inherently long relaxation time of the (liquid) sample, (and hence low signal-to-noise ratio (SNR)) causes unnecessarily long repetition time. Furthermore, the T_2_-relaxation measurement (in one-dimensional) which is frequently reported in NMR relaxometry experiments has limited number of spaces (e.g., healthy/non-healthy)^[20,21]^.

In this work, we present a new class of methodology for rapid and accurate object classification using PoCT NMR relaxometry with the aid of machine learning (Fig. 1). We demonstrated (using various edible oils as proof-of-concept) that the sensitivity (‘true positive rate’) and specificity (‘true negative rate’) of the classification is substantially improved using higher order of (pseudo)-dimensionality (e.g., 2D or multidimensional). Further, by leveraging on the advances in machine learning techniques (e.g., pre-trained dataset) the detection time was sped up (in minutes) as compared to conventional 2D or multidimensional NMR (>hours), without resorting to using Ultrafast NMR^[28]^. This methodology (termed as Clustering NMR) is extremely useful for rapid and accurate classification of objects (in less than a minute) using the low-field NMR at point-of-need.

**Figure 1:**
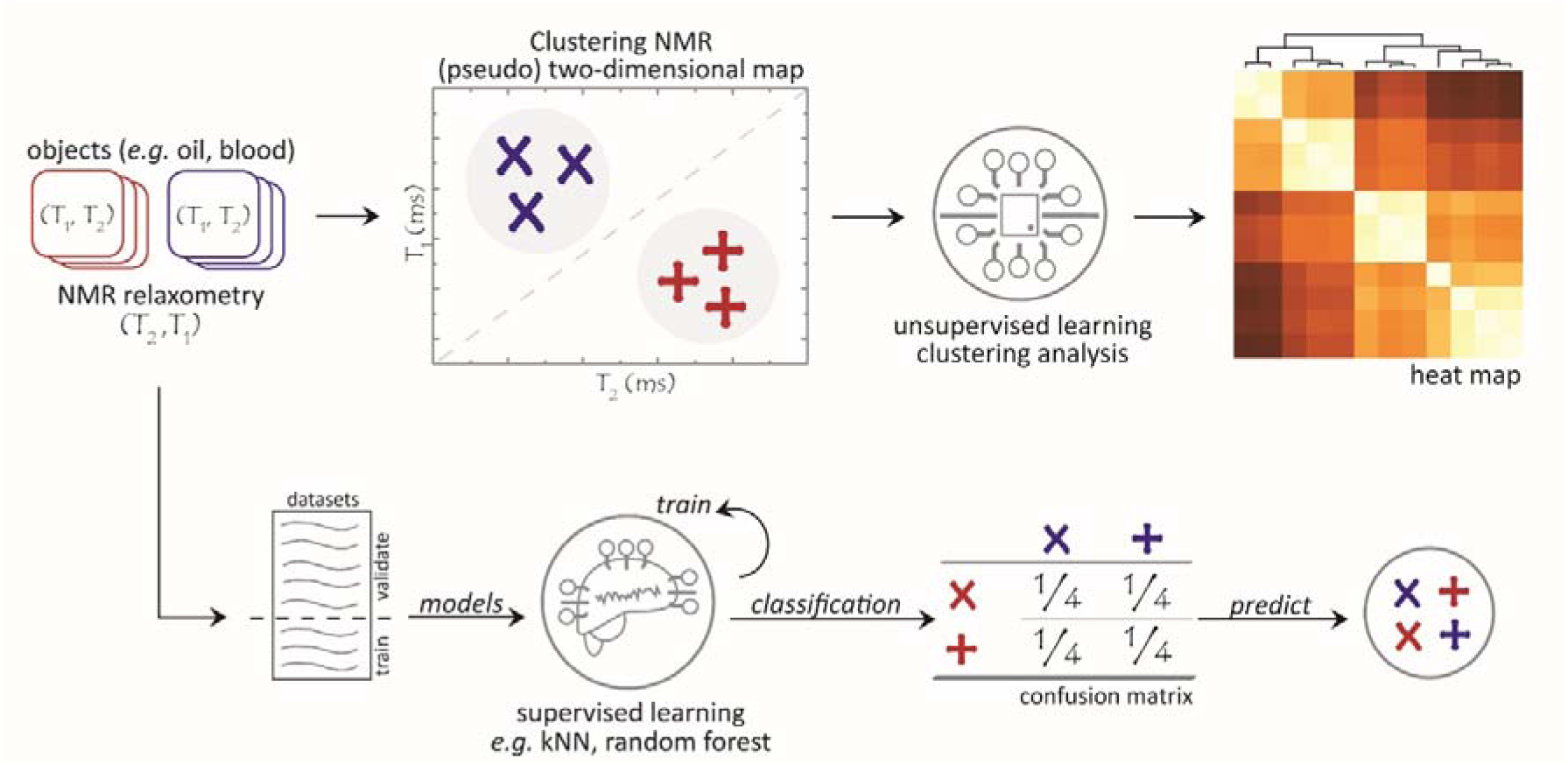
Conceptualization of the (pseudo) two-dimensional mapping using the Clustering NMR method proposed in this work. A pair of (T_1_, T_2_) relaxation time for each objects (e.g., edible oils, blood) were measured using micro NMR relaxometry system. A (pseudo) two-dimensional map is constructed with (T_2_, T_1_) relaxation time with a (X, Y) scatter plot (Fig. 2c), where the object clustering became obvious in comparison to its’ one-dimensional counterparts (i.e, T_1_ relaxation or T_2_ relaxation). The efficacies of Clustering NMR method were validated using both the supervised and unsupervised learning methods. The relationship between each objects is established using (unsupervised) clustering analysis methods (e.g., tree classfication, hierarchical clustering) and its’ quantitative linkage (e.g., inter/intra cluster similarity) of each objects which is depicted on a dendogram with a heat map (details in Supp. Figs. 2-3). Supervised learning techniques (e.g., kNN, random forest, logistic regression) were used to train the classification of objects and the best trained model is subsequently chosen to predict the object classification. (e.g., oils content, infection/non-infection).

## Methods

NMR measurement and detection. The relaxometry measurements (T_1_ relaxation, T_2_ relaxation) were carried out on four group edible oils (i.e., peanut, olive, sunflower, corn) labelled as (A, B, C, D), respectively (Figs. 2a-b). In order to avoid bias, more than one different manufacturers were used for the same oil (with the exception of corn oil) and the detail on fat compositions were presented in Supp. Fig. 1. (A, A’, B, B’, C, C’, C’’, D) were the variants of the same oil from various manufacturers. The manufacturer labelling indicated 100% of oil contents (no mixture of oils). The edible oils were cooking oils bought locally in Braga, Portugal. No further alteration was made before the NMR measurements.

**Figure 2:**
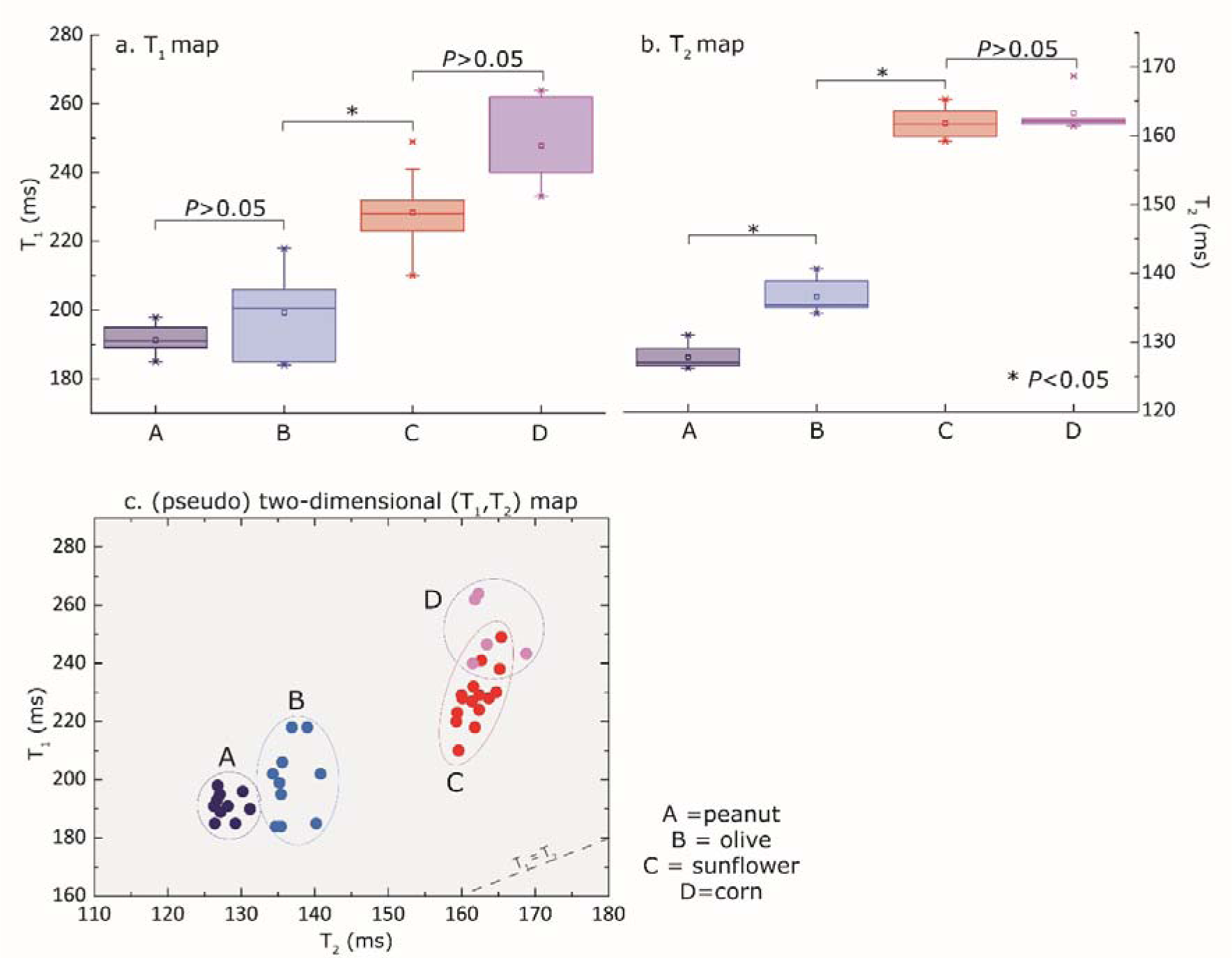
NMR measurements and (pseudo) two-dimensional mapping with Clustering NMR approach. T_1_ and T_2_ relaxation times were carried on various edible oils (i.e., peanut, olive, sunflower, corn), with the label of (A, B, C, D) respectively using micro NMR relaxometry. One-dimensional mapping with (a) T_1_ relaxation time, (b) T_2_ relaxation time, and (c) (pseudo) two-dimensional mapping using a pair of (T_1_, T_2_) relaxation times. NMR measurements were carried out on each edible oils in quintuplicate manner with a total of 40 points (datasets). The clustering circles were drawn for eye-balling purposes. Details of the oils (e.g., manufacturers, fat compositions) were presented in Supp. Fig. 1. The box plots represent 25% and 75% quantile of the entire measurements. The diaganol line (T_1_=T_2_) represents the border limit where it is physically non-measurable. Two tailed Student’s T-test was used to calculate the *P*-value.

NMR measurements were carried out (in single blinded manner) on each oils in quintuplicate manner (i.e., five repeated times) with a total of 40 points for all the samples. Details on NMR parameter are reported in Supplementary Methods. Clustering NMR method uses a pair of (T_2_, T_1_) relaxation time for each objects (e.g., edible oils, blood) to construct a (pseudo) two-dimensional map (Fig. 2c). The pseudo two-dimensional map can be used a referencing map (control).

Machine learning learning algorithm and workflows. Using a statistical programming languages (e.g., *R* or Orange 3.1.2), the raw datasets can be processed using supervised and unsupervised learning techniques. The machine learning algorithms were written and runs on a personal laptop (Intel Core Pentium i7 CPU @ 2.70GHz, 8.00 GB RAM). Once the model in machine learning is built, all the tasks run simultaneously and completes typically in less than 1 minute.

Using unsupervised learning, the relationship between each objects were rapidly constructed using clustering analysis (e.g., tree classification, hierarchical clustering) and its’ quantitative linkages (e.g., inter/intra cluster similarity) were shown on a dendogram and a heat map (Fig. 3). Supervised learning models (i.e., neural network, kNN, logistic regression, naïve Bayes, and random forest) can be used to train the datasets and the best model with the highest accuracy can be chosen to predict the object classification (e.g., oil classification, infection/non-infection) using pre-trained datasets (Fig. 4 and Fig. 5).

**Figure 3:**
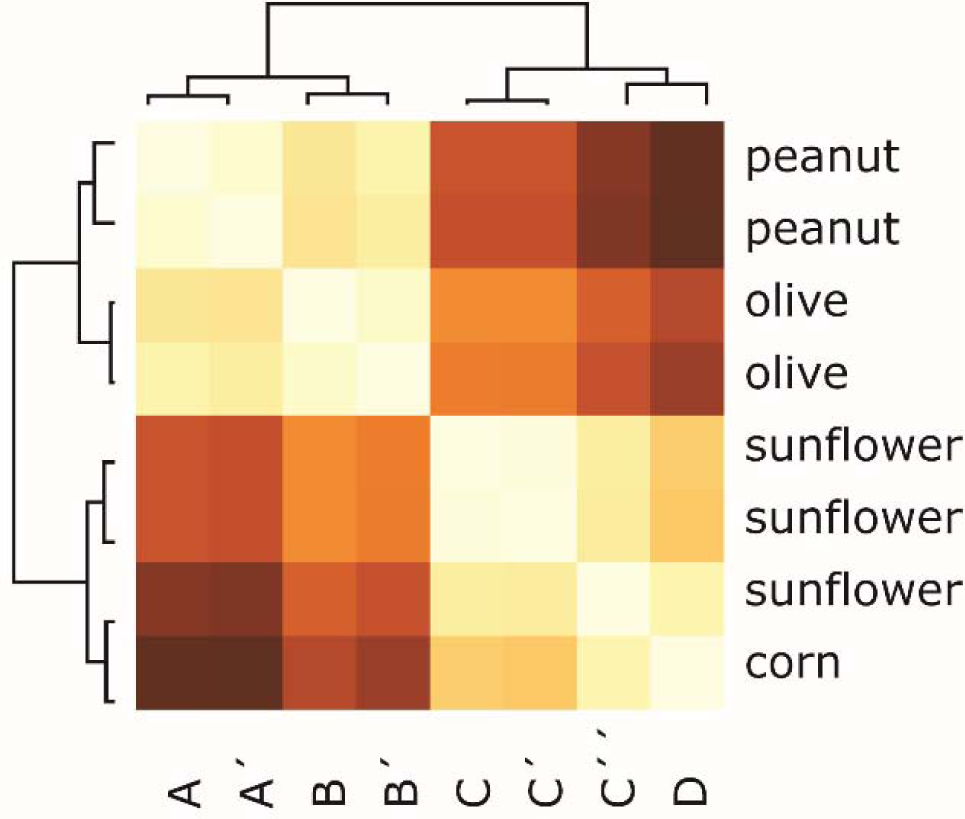
Unsupervised learning techniques (e.g., hierarchical clustering, tree classification) can be used for clustering analysis. This hierarchical clustering was constructed based on Euclidean distance (between T_1_ relaxation and T_2_ relaxation) and its’ quantitative linkages (e.g., inter/intra cluster similarity) shown in a heat map. (A, A’, B, B’, C, C’, C’’, D) were the variants of the same oil content taken from different manufacturers. Tree classification method is shown for comparison (Supp. Fig. 2).

**Figure 4:**
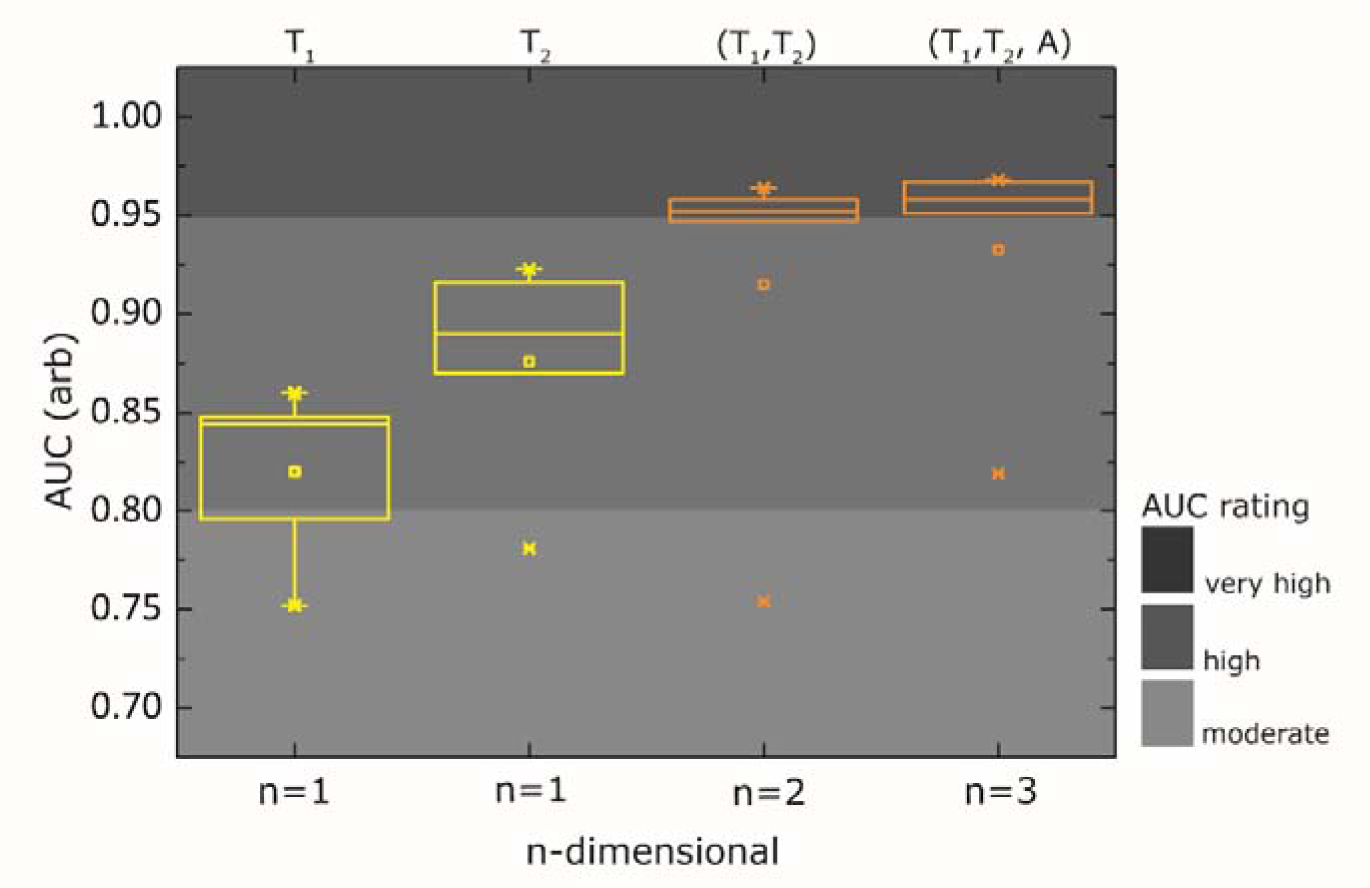
The Area Under Curve (AUC) plot as evaluated by Receiver Operating Characteristic (ROC) of various supervised models (i.e., kNN, random forest, neural network, naïve bayes, logistic regression) evaluated using target variables of (a) one-dimensional (T_1_-relaxation), (b) one-dimensional (T_2_-relaxation), (c) two-dimensional (T_1_-relaxation, T_2_-relaxation), and (d) three-dimensional (T_1_-relaxation, T_2_-relaxation, A-ratio) using leave-one-out training method. Other details can be found in Table 1.

## Results

Each edible oils (i.e., peanut, olive, sunflower, corn) were assigned to its’ respective label (A, B, C, D) following the blinded NMR measurements. As depicted in the one-dimensional map, each of oil contents has a specific T_1_ relaxation and T_2_ relaxation characteristic reading (Figs. 2a-b). The means for T_1_ relaxation time were (191.3, 199.3, 228.4, 247.8) ms and means for T_2_ relaxation time were (127.9, 136.8, 162, 163) ms for (A, B, C, D), respectively.

**Table 1:** The sensitivity and specificity of the various supervised models evaluated using target variable/s of (a) one-dimensional (T_1_-relaxation), (b) one-dimensional (T_2_-relaxation), (c) two-dimensional (T_1_-relaxation, T_2_-relaxation), and (d) three-dimensional (T_1_-relaxation, T_2_-relaxation, A-ratio) from leave-one-out training method. The synonyms used were: T_1_-relaxation/T_2_-relaxation (A-ratio), area under the curve (AUC), classification accuracy (CA), F1 score – the balance between precision and recall, Precision – how many selected items were relevant, Recall - how many relevant items are selected. The training method using cross validation of k=5 was also evaluated for comparison (Supp. Fig. 4).

|  | Method | AUC | CA | F1 | Precision | Recall |
| --- | --- | --- | --- | --- | --- | --- |
| a) | kNN | 0.848 | 0.575 | 0.563 | 0.561 | 0.575 |
|  | Random Forest | 0.860 | 0.600 | 0.598 | 0.598 | 0.600 |
|  | Neural Network | 0.844 | 0.650 | 0.626 | 0.633 | 0.650 |
|  | Naive Bayes | 0.796 | 0.625 | 0.575 | 0.542 | 0.625 |
|  | Logistic Regression | 0.752 | 0.425 | 0.333 | 0.288 | 0.425 |
|  | average | 0.820 | 0.575 | 0.539 | 0.524 | 0.575 |
| b) | kNN | 0.923 | 0.750 | 0.750 | 0.750 | 0.750 |
|  | Random Forest | 0.916 | 0.750 | 0.750 | 0.750 | 0.750 |
|  | Neural Network | 0.870 | 0.825 | 0.771 | 0.740 | 0.825 |
|  | Naive Bayes | 0.890 | 0.875 | 0.821 | 0.781 | 0.875 |
|  | Logistic Regression | 0.781 | 0.625 | 0.481 | 0.392 | 0.625 |
|  | average | 0.876 | 0.765 | 0.715 | 0.683 | 0.765 |
| c) | kNN | 0.958 | 0.775 | 0.773 | 0.792 | 0.775 |
|  | Random Forest | 0.947 | 0.850 | 0.828 | 0.827 | 0.850 |
|  | Neural Network | 0.964 | 0.850 | 0.840 | 0.850 | 0.850 |
|  | Naive Bayes | 0.952 | 0.775 | 0.766 | 0.758 | 0.775 |
|  | Logistic Regression | 0.754 | 0.600 | 0.521 | 0.476 | 0.600 |
|  | average | 0.915 | 0.770 | 0.746 | 0.741 | 0.770 |
| d) | kNN | 0.958 | 0.775 | 0.773 | 0.792 | 0.775 |
|  | Random Forest | 0.968 | 0.875 | 0.873 | 0.874 | 0.875 |
|  | Neural Network | 0.951 | 0.800 | 0.793 | 0.809 | 0.800 |
|  | Naive Bayes | 0.967 | 0.825 | 0.830 | 0.836 | 0.825 |
|  | Logistic Regression | 0.819 | 0.625 | 0.568 | 0.572 | 0.625 |
|  | average | 0.933 | 0.780 | 0.767 | 0.777 | 0.780 |

The spread of the readings were, however, substantially large making objects (A and B) and objects (C and D) inseparable in the T_1_ relaxation dimension (P>0.05) (Fig. 2a). Further in the T_2_ relaxation dimension, the objects (C and D) were also inseparable (Fig. 2b). The undesirable spread causes (similarly to spectral) cluster overlapping and hence making classification difficult (if not impossible). One straightforward solution is to increase the SNR (e.g., increasing the number scans) or/and increase the number of samplings, which unfortunately, came at the expenses of acquisition time. In addition, the relaxation time of liquid sample is inherently long. On the other hand, using the Clustering NMR method (as proposed in this work), one can leverages on the combined characteristic of (T_1_, T_2_) relaxation times of the oil contents. It forms (visibly) unique and specific cluster based on the oil contents (‘molecular fingerprint’) in (pseudo) two-dimensional map (Fig. 2c). With the minor exception of corn oil (which partially overlapped with sunflower oils), which could be due to possible adulteration or factory processes. Upon further investigation, we found that this artifact can be removed with higher SNR.

Interestingly, unsupervised techniques based clustering analysis (e.g., hierarchical clustering (HC), tree-based classification, and k-means) can be performed in conveniently using (open-source code) user friendly third party software (e.g., *R*, or Orange 3.1.2). A front-end statistical programming language allows the clustering analysis (once compiled), can be executed in the next occasion. The HC analysis successfully separated the (peanut and olive) cluster from the (sunflower and corn) cluster, and subsequently split between themselves (Fig. 3). The HC was constructed based on Euclidean distance (between T_1_ relaxation and T_2_ relaxation) and its’ quantitative linkages (e.g., inter/intra cluster similarity) were shown in a heat map. The HC methods also confirmed the oil variants (A, A’, B, B’, C, C’, C’’, D) based on its’ respective manufacturer. Similarly, the Chemometric approach^[29]^ based on fat compositions (Supp. Fig. 2) and tree-based classification technique based on the T_1_-relaxation cutoff and T_2_-relaxation cutoff criterion (Supp. Fig. 3) appear to be in good agreement (qualitatively) with the HC classification using Euclidian distance of T_1_ relaxation and T_2_ relaxation obtained with NMR experimentally. It is worth noting, however, that the figures (i.e., fat compositions) given by the manufacturers are for references (and not for scientific) purposes. The clustering analysis models despite using various differential clustering criterions (e.g., Euclidean distance, fat compositions, relaxation cutoff) were in agreement with our observation (Clustering NMR, Fig. 2c). This demonstrated the robustness of Clustering NMR method, which can be validated using unsupervised techniques.

In order to evaluate the classification accuracy on the quantitative basis, various supervised learning models (i.e., kNN, random forest, neural network, naïve Bayes, and logistic regression) were used to train, validate and predict the datasets. The Area Under Curve (AUC) as evaluated with Receiver Operating Characteristic (ROC) were on average (0.820, 0.876, 0.915, 0.933) with (one-dimensional (T_1_-relaxation), one-dimensional (T_2_-relaxation), two-dimensional (T_1_-relaxation, T_2_-relaxation), and three-dimensional (T_1_-relaxation, T_2_-relaxation, A-ratio)), respectively, using the leave-one-out training method (Fig. 4). A-ratio is the ratio between T_1_-relaxation and T_2_ -relaxation. Similar conclusions were observed using cross validation method (e.g., k=5) (details in Supp. Table 1). This confirmed that the sensitivity and specificity of the proposed Clustering NMR method has substantially improved at the higher order of (pseudo)-dimensionality (e.g., 2D or multidimensional) over low dimensionality (e.g., n=1). With the (minor) exception of logistic regression, all the supervised models performed reasonably well (AUC>0.80) (Table 1). Furthermore, all the machine learning tasks run simultaneously and computational time taken were typically in less than 1 minute (in this work).

## Discussion

The proposed Clustering NMR method works on the rational that accumulative characteristic of each dimensionality would forms a specific and unique signature (‘molecular fingerprint’). This is the same concept which borrowed from the data mining^[30]^. Fortunately, the characteristic of (T_1_, T_2_) relaxation times in the relaxometry is rather specific and prominent, and as the results suggested, an optimal n=2 to 3 of dimensionality are essential to attain a high AUC (Fig. 4). This is much smaller than n>thousands when the effect of ‘curse of dimensionality’ ^[31]^ became prominent. With the recent advances in machine learning, however, its’ becoming computationally cheaper (e.g., shorter analysis time) to calculate a big dataset. The computational time reported in this analysis (less than one minute) much shorter than a conventional two- or multidimensional NMR (>hours), without resorting to the use of Ultrafast NMR.

Two- or multidimensional relaxometry experiments (e.g., T_1_-T_2_ correlation spectroscopy), however, may provides much more information (e.g., cross peaks) but are far more time consuming than that of Clustering NMR method. One way to speed up acquisition time is to employ the use of gradient fields (e.g., Ultrafast NMR^[28]^, continuous spatial encoding^[32]^) which require modification to the radio-frequency probe. Machine learning in the form of dimension reductionist (e.g., principal component analysis (PCA), partial least squares (PLS)) have also been used to reduce the dimensionality in multidimensional spectroscopy (e.g., NMR metabolomics ^[19,33,34]^). A recent deep learning assistive NMR spectroscopy^[18]^, which signals reconstructing were demonstrated. We summarized and compared Clustering NMR method with the state-of-the-art methodologies in a SWOT-like analysis (Table 2).

**Table 2:** State-of-the-art (with/without) machine learning assistive NMR works in comparison to the current work (Clustering NMR).

| Authors/Year | NMR | Applications | n-Dimensional | Machine Learning | Advantageous | Weakness |
| --- | --- | --- | --- | --- | --- | --- |
| Wishart (2008)<sup>[19]</sup>, Karaman (2015)<sup>[34]</sup>, Rocha (2018)<sup>[33]</sup> | spectroscopy | metabolomics | 2D | PCA/PLA | informative | slow |
| Frydmann (2014)<sup>[28]</sup> | spectroscopy | ultrafast NMR | 2D | no | rapid | gradient field |
| Qu (2019)<sup>[18]</sup> | spectroscopy | generic | n-dimensional | deep learning | speed up | information lose? |
| Haun (2010)<sup>[38]</sup>, Haun<sup>[20]</sup> (2011), Liong (2013)<sup>[35]</sup>, Peng (2014)<sup>[21]</sup>, Neely (2016)<sup>[39]</sup>, Robinson (2017)<sup>[40]</sup> | relaxometry | medical diagnosis | 1D | no | rapid, PoCT | low specificity and sensitivity, missing out cross peaks?! |
| Robinson (2014)<sup>[41]</sup>, Ok (2016)<sup>[42]</sup> | relaxometry | food science | 1D | no | rapid, PoCT |  |
| Santos (2016)<sup>[43]</sup>, Zhu (2016)<sup>[38]</sup> | relaxometry | food science | 1D | PCA/PLA | rapid, PoCT |  |
| Xu (2014)<sup>[44]</sup>, Rudszuck (2019)<sup>[27]</sup> | relaxometry | food science | 2D | no | PoCT, high specificity and sensitivity | slow |
| Hurlimann (2002)<sup>[45]</sup> | relaxometry | oil-gas exploration | 1D, 2D | no | *in situ*, high specificity and sensitivity | slow |
| Lewis (2013)<sup>[46]</sup> | relaxometry | oil-gas exploration | 2D | no |  | slow |
| Birdwell (2015)<sup>[47]</sup> | relaxometry | oil-gas exploration | 2D | PCA/PLA |  | slow |
| Clustering NMR (2020) | relaxometry-clustering | generic | (pseudo) n-dimensional | clustering analysis, supervised model | rapid, PoCT, high specificity and sensitivity | missing out cross peaks? |

In conclusion, this proposed methodology, termed as Clustering NMR is extremely powerful for rapid and accurate classification of objects using the low-field NMR. This methodology is highly distruptive to the low-field NMR applications, in particularly, the recent reported PoCT medical diagnostic. These include the immuno-magnetic labelled detection (e.g., tumour cells ^[14,20]^, tuberculosis^[35]^ and magneto-DNA detection of bacteria^[36]^) and the label-free detection of various pathological states (e.g., blood oxygenation^[15]^/oxidation level^[10]^ and malaria screening^[21,22,37]^). Interestingly, with the recent advances on machine learning technique, it has become remarkably efficient that a large data run in almost in ‘real-time mode’, which open-up opportunity to combine real-time NMR (or MRI) with machine learning simultaneously.

## Supplementary Methods

NMR setup and parameters. The ^1^H magnetic resonance measurements of edible oils were carried out at the resonance frequency of 21.67 MHz using a portable permanent magnet (Metrolab Instruments, Switzerland), B_o_=0.5T using a benchtop-type console (Kea Magritek, New Zealand). A temperature controller was set to maintain the measurement chamber at 30°C. The T_1_ relaxation and T_2_ relaxation pulse sequences were set at standard inversion recovery, followed by Carr-Purcell-Meiboom-Gill (CPMG) train pulses, respectively. The experimental parameters used were echo time=200 µs, number of echoes=2000 and signal averaging=4. A recycle delay of 4s was set between each experiment to provide sufficiently long time to allow all the molecular spins to return to thermal equilibrium.

## Statistical methods

Two tailed Student’s T-test was used to calculate the *P*-value.

## Data availability statement

The machine learning algorithms and raw NMR datasets are available upon reasonably request at.

## Acknowledgement

This research was supported by the International Iberian Nanotechnology Laboratory (INL) Start Up and INL Seed Grant. W.K. Peng would like to personally thank internship student Daniele Paesani and Dr Murali Kumarasamy, who carried out the NMR measurements in blinded manner and for assisting other laboratory works.

## Author Contribution

W.K.P conceived the original idea, wrote the paper, designed the protocols, and built the entire hardware setup.

## Additional information

Supplementary information is available for this paper.

